# Evaluating Human Autosomal Loci for Sexually Antagonistic Viability Selection in Two Large Biobanks

**DOI:** 10.1101/2020.03.26.009670

**Authors:** Katja R. Kasimatis, Abin Abraham, Peter L. Ralph, Andrew D. Kern, John A. Capra, Patrick C. Phillips

## Abstract

Sex and sexual differentiation are ubiquitous across the tree of life. Because females and males often have substantially different functional requirements, we expect selection to differ between the sexes. Recent studies in diverse species, including humans, suggest sexually antagonistic viability selection creates allele frequency differences between the sexes at many different loci. However, theory and population-level simulations indicate that sex-specific differences in viability would need to be very extreme in order to produce and maintain reported levels of between-sex allelic differentiation. We address this paradox between theoretical predictions and empirical observations by evaluating evidence for sexually antagonistic viability selection on autosomal loci in humans using the largest cohort to date (UK Biobank, n=438,427) along with a second large, independent cohort (BioVU, n=93,864). We performed association tests between genetically ascertained sex and genotypes. Although we found dozens of genome-wide significant associations, none replicated across samples. Moreover, closer inspection revealed that all associations are likely due to cross-hybridization with sex chromosome regions during genotyping. We report loci with potential for mis-hybridization found on commonly used genotyping platforms that should be carefully considered in future genetic studies of sex-specific differences. Despite being well-powered to detect allele frequency differences of up to 0.8% between the sexes, we do not detect evidence for this signature of sexually antagonistic viability selection on autosomal variation. These findings suggest a lack of strong ongoing sexually antagonistic viability selection acting on single locus autosomal variation in humans.

## INTRODUCTION

Understanding the relationship between genotype and sexually dimorphic phenotypes, and how selection shapes this relationship, is fundamental to understanding sex-specific responses in aging (Archer *et al.* 2018), fertility (Farquhar *et al.* 2019), disease susceptibility (Morrow 2015; Ferretti *et al.* 2018; Dumitrescu *et al.* 2019), and treatment (Khramtsova *et al.* 2018). Sexual dimorphism is common across a range of plant and animal taxa (Rowe *et al.* 2018; Deegan and Engel 2019). Differences in optimal fitness values between the sexes may result in sexually antagonistic selection (Arnqvist and Rowe 2005) – i.e., selection on autosomal variants that affect fitness in different directions for each sex. Surveys of natural selection suggest that the repeated evolution of sexual dimorphism is commonly associated with sexually antagonistic selection (Cox and Calsbeek 2009). Yet, we still lack an understanding of how this process shapes genomic variation within and between species. A major obstacle in assessing the genomic consequences of sexually antagonistic selection is that most of the hypothesized genomic signatures are not unique to this mode of selection. However, when the alleles at a single locus have opposite effects on viability between the sexes, namely intralocus sexual conflict (Rice and Chippindale 2001), then sexually antagonistic viability selection is hypothesized to favor different alleles in each sex. This process is predicted to generate allele frequency differences between the sexes among adults (Mank 2017; Kasimatis *et al.* 2017).

Recent research has looked for this signature of selection by identifying alleles with high male-female *F*_*ST*_ (Cheng and Kirkpatrick 2016; Kasimatis *et al.* 2017; 2019), a normalized measure of allele frequency difference. Such studies across a range of taxa have suggested that potentially hundreds of autosomal loci are subject to ongoing sexually antagonistic selection with many differentiated loci having male-female divergence values of at least 10% (Lucotte *et al.* 2016; Flanagan and Jones 2017; Wright *et al.* 2018; Dutoit *et al.* 2018; Bissegger *et al.* 2019), and some reaching even as high as 45% (Vaux *et al.* 2019). These results are surprising because the production and maintenance of such large male-female differences on autosomes requires strong, ongoing selection to overcome the homogenization of genotypes during meiotic segregation every generation (Cheng and Kirkpatrick 2016; Kasimatis *et al.* 2019). Theory suggests that a male-female *F*_*ST*_ value of 1% requires at least a 33% viability cost per sex per generation (Kasimatis *et al.* 2019). Given the high sex-specific viability cost, factors such as population structure, sampling variance due to small sample sizes, or bioinformatic artifacts may contribute to the high divergence values observed (Kasimatis *et al.* 2019). Of particular concern are the small sample sizes (15 to 100 individuals) used by many previous studies. Detecting the level of allelic differentiation expected at sexually antagonistic loci with moderate sex-specific mortality (≤10% per sex) requires substantially larger sample sizes and accounting for other possible confounding effects, such as population structure (Kasimatis *et al.* 2019). Indeed, a meta-analysis of 51 studies that included more than 100,000 European-ancestry individuals did not find any common variants associated with sex ratio (Boraska *et al.* 2012).

We aim to reconcile empirical observations with theoretical predictions using a robust statistical framework to identify intralocus sexually antagonistic viability selection in the largest human cohort to date. We use two large-scale biobanks, the UK Biobank and the Vanderbilt Biobank (BioVU) to analyze >500,000 human genomes for signals of male-female divergence driven by sexually antagonistic selection. We rigorously control for population stratification and potential molecular and informatic artifacts. Compared to previous studies examining sexual antagonism, these datasets significantly improve our statistical power to detect allele frequency differences among the sexes by providing the largest available sample sizes to date – several orders greater than previous studies in humans (Lucotte *et al.* 2016; Cheng and Kirkpatrick 2016) and non-model taxa (Lucotte *et al.* 2016; Flanagan and Jones 2017; Wright *et al.* 2018; Dutoit *et al.* 2018; Bissegger *et al.* 2019; Vaux *et al.* 2019). Our association framework differs from traditional association studies as genetic sex is the phenotype of interest and the mechanism generating a true effect would be sex-specific viability. After controlling for multiple confounders, we are unable to detect evidence for ongoing sexually antagonistic viability selection at individual autosomal loci.

## MATERIALS AND METHODS

### Genotyping and quality control in BioVU

The DNA biobank at Vanderbilt University, consists of DNA extracted from blood collected during routine clinical testing. For 93,864 individuals, GWAS-level genotyping was performed using the Illumina MEGA-Ex chip which includes >2 million common and rare variants before imputation. We obtained genotyped data in PLINK format from the Vanderbilt sequencing core after the following quality control steps: excluding either samples or variants with ≥5% missingness, mismatched identifiers as detected by identity by descent checks, and non-concordance between reported gender and genetically determined sex. Overlapping variants with 1000 Genomes demonstrated ≥99.98% variant call concordance using HapMap sample aliquots. Using PLINKv1.90b3s (Chang *et al.* 2015), we additionally performed the following quality control steps. We first confirm that duplicate samples and those with high missing rate (≥5%) are not present and exclude samples with high heterozygosity on autosomes (>3 S.D. from observed data), or high relatedness (%IBD ≥ 0.2). Next, we removed duplicated variants and variants with high missing rate (≥5%) or significantly different missing rate between cases and controls (p < 0.00001, Fisher’s Exact test). We then included only samples with a self or third party reported race as ‘white’ and variants with minor allele frequency >0.01. This additional quality control resulted in a final European-ancestry dataset of 61,760 samples (34,269 females and 27,491 males) and 1,763,607 variants. We calculated the top 12 principal components on this cohort. We imputed variants that reached nominal or genome-wide statistical significance (P < 5E-8) in the UK Biobank data but were not genotyped in the BioVU cohort. These variants were imputed using the Michigan Imputation Server (v1.2.4) (Das *et al.* 2016) using the HRC (Version r1.1 2016) reference panel and retaining variants with R^2^ > 0.3. Imputed allele dosages were converted to hard calls using PLINK/2.00-alpha2 (Chang *et al.* 2015) and filtered to exclude variants with minor allele frequency <1% and genotyping rate <95%. All PLINK code is available on the GitHub repository https://github.com/abraham-abin13/sexually_antagonistic_sel.git.

### Genotyping and quality control in the UK Biobank

The UK Biobank is an international health resource with data from approximately 500,000 participants. Genotyping and quality control procedures have previously been described in detail by Bycroft et al. (2018). Briefly, two arrays – the UK Biobank Axiom Array (n = 438,427 participants) and the UK BiLEVE Axiom Array (n = 49,950 participants) – were used to genotype participants (71 bp oligos). Quality control procedures carried out before the data were released, included: removal of participants with excess heterozygosity or missingness, removal of markers with batch, plate, array, or sex effects, and removal of markers with discordance across control replications (Bycroft *et al.* 2018). The removal of sex effects, namely allele frequency differences between females and males at a given marker, does not preclude our analysis as the conservative threshold (P < 10E-12) removed only eight markers and the sex differences at these markers were due to technical artifacts, such as the probe sequence mapping to the Y chromosome (C. Bycroft pers. comm.). The released genotype data contains 805,462 markers from 488,377 participants (Field IDs 22100-22124). Additionally, the genetic sex (Field ID 22001), year of birth (Field ID 34), date of assessment (Field ID 53), and assessment center (Field ID 54) were requested for each participant. The top 40 genetic principal components (Field ID 22009) were previously calculated using fastPCA (Bycroft *et al.* 2018).

Using PLINKv1.90b3s (Chang *et al.* 2015), we additionally performed the following quality control steps. We excluded samples with high missing rate (≥5%) and high heterozygosity on autosomes (>3 S.D. from observed data). Next, we pruned markers in linkage-disequilibrium (window size = 50kb, step rate = 5, r^2^ threshold = 0.2). Finally, we removed variants with significantly different missing rate between cases and controls (P < 0.00001, Fisher’s Exact test). We included only variants with minor allele frequency > 0.01 to exclude inaccurate calls made for low frequency alleles (Wright *et al.* 2019; Weedon *et al.* 2019). This additional quality control resulted in a final dataset of 488,291 samples (264,813 females and 223,478 males) and 653,632 variants. A binomial test was used to test for a lack of minor allele homozygotes relative to that expected under HWE (this is conservative, because most human population dynamics is expected to lead to an excess of homozygosity). All PLINK code is available on the GitHub repository https://github.com/abraham-abin13/sexually_antagonistic_sel.git

Imputed genotype and phased haplotype values were used to compare significant loci in the BioVU cohort, which were not directly genotyped in the UK BIOBANK arrays. Again, imputation was completed prior to the data release using the Haplotype Reference Consortium and UK10K haplotype resource. The imputation methods are described in detail in Bycroft et al. (2018). Imputed allele dosages were converted to hard calls using PLINK/2.00-alpha2 (Chang *et al.* 2015).

### Genome-wide association for an individual’s sex

We performed a GWAS in UK Biobank and BioVU separately using a logistic regression testing the association between an individual’s sex (binary variable, concordant with their genetic sex) and the effect allele, defined as the minor allele by PLINKv1.90b3s (Chang *et al.* 2015), using an additive model. For BioVU analysis, we controlled for genetic ancestry using 12 genetic principal components and included year of birth as a covariate. For the UK Biobank analysis, we again controlled for genetic ancestry using 12 genetic principal components, along with age at assessment and UK Biobank sampling center as covariates. All genome wide association tests were done using PLINKv1.90b3s (Chang *et al.* 2015). We focused our analyses on the autosomes, where genomic divergence between the sexes is not confounded by sex chromosome processes. During our quality control steps before association testing, we did not remove variants based on deviations from Hardy Weinberg Equilibrium (HWE) since theory indicates that sex-specific selection can violate the assumptions of HWE (Kasimatis *et al.* 2019).

### Resampling of sex and generating a null distribution

To determine if p-values were well calibrated (i.e., uniformly distributed on [0,1]) at non-associated variants, we performed a permutation analysis to calculate the distribution of p-values within the UK Biobank cohort. We resampled genetic sex 100 times per chromosome to generate a set of random associations between genotype and this phenotype. We then reran the logistic regression, again including 12 genetic principal components, age, and sampling center as covariates, for only those variants that had a p-value < 0.01 in the original association analysis (n = 8,868 SNPS). These analyses generated a distribution of 100 p-values at each variant. Permuted p-values were uniformly distributed (Supplementary Figure 1A), even when the values were small (Supplementary Figure 1B), indicating the p-values for this association analysis were well-calibrated and therefore a genome-wide Bonferroni significance threshold of P < 5E-8 was appropriate. All R and PLINK code are available on the GitHub repository https://github.com/abraham-abin13/sexually_antagonistic_sel.git.

### Identifying SNPs with sequence similarity to sex chromosomes

Incorrectly mapped sex-chromosome variants to an autosomal region can result in statistically significant GWAS hits for an individual’s sex due to the different effects on allele counts between females and males. We used BLAT (Kent 2002) with default parameters (stepSize=5, repMatch=2253, minScore=20, minIdentity=0) to identify sequence similarity between the probe sequences used on the genotyping array and sex chromosome regions. The MEGA-Ex array probe sequences used to genotyped the BioVU cohort were obtained directly from Illumina. Probes sequence for the UK Axiom Biobank array (Resource 149601) and UK BiLEVE array (Resource 149600) were download from https://biobank.ctsu.ox.ac.uk/crystal/label.cgi?id=263. MEGA-Ex probes are 50 base pair sequences adjacent to the variant being tested; MEGA-Ex uses single base extension to detect the variant allele. UK Biobank array probes are 71 base pairs long with the variant being genotyped located in the middle. BLAT hits to the X or Y chromosome were further filtered to identify regions likely to cross-hybridize by requiring at least 40 base pair overlap, sequence similarity ≥90%, and that the matching sequence overlaps (UK Biobank arrays) or flanks (MEGA-Ex array) the variant being tested. Similar criteria were used in a previous a study that reported cross-hybridization on the Illumina Infinium HumanMethylation27K microarray platform (Chen *et al.* 2012). Next, we identified the best BLAT hit to a sex chromosome for each probe sequence by selecting the hit with the highest BLAT score, which accounts for match length and sequence similarity. For this step, we considered the UK Axiom and UK BiLEVE array together thus selecting the probe sequence with the highest BLAT score from one of the two arrays per variant tested in the GWAS. In the BioVU (MEGA-Ex array) and UK Biobank arrays, 83,083 out of 798,051 and 128,090 out of 620,040 autosomal probes had at least one BLAT match (BLAT score ≥ 20) to a sex chromosome region. To further focus on sequence similarity with potential to cause genotyping error, we identified sex chromosome matches with the following criteria (Chen *et al.* 2012): 1) ≥40 base pairs in length, 2) ≥90% sequence similarity, and 3) overlap between the match and the variant being genotyped.

### Power Analysis

We conducted a power analysis to determine how the minimum allelic divergence between the sexes that could be detected within the BioVU and UK Biobank cohorts (Supplemental File 9). Specifically, we determined the probability that we would reject the null observation that the population frequency of each allele is equal at a p-value threshold of P = 1E-8. Suppose we have N males and M females, and the allele frequencies in the two groups are P and Q. Since the cohort sample sizes are large, if the population frequencies are *p* and *q*, then P ∼ Normal(*p, p* (1-*p*) / 2N) and Q ∼ Normal(*q, q* (1-*q*) / 2M). The difference in population allele frequencies is then given by P - Q ∼ Normal(*p* - *q, p* (1-*p*) / 2N + *q* (1-*q*) / 2M). The variance is maximized when *p* = *q* = 1/2, so is at most the variance in the population is: V = (1/N + 1/M)/8. The two-sided p-value for P-Q being nonzero will be below 1E-8 if |P-Q| is larger than z(0.5e-8) *sqrt(V), where z(*p*) is the p-th quantile for the standard Normal distribution. Even, then, an allele with |P-Q| = z(0.5E-8) * sqrt(V) will only have a two-sided p-value half the time; alleles must by slightly farther apart (by z(0.025) * sqrt(V)) to have a 95% probability that statistical noise does not put them above the p=1e-8 threshold. Therefore, we will have 95% power to detect any SNP with true |p-q| > (z(0.5E-8) + z(0.025)) * sqrt(V).

### Data Accessibility

All the data generated from this study (Supplemental Files 1-9) were deposited in the figshare repository https://figshare.com/s/e863ea11cc9dab30c1b9 to be made public upon publication. All the code generated for this study were deposited in a GitHub repository to be made public upon publication.

## RESULTS

Throughout this paper when we refer to an individual’s sex, we are referencing that individual’s sex chromosome composition as estimated in each biobank dataset and binarized (i.e., metadata reports each individual as XY or XX, although the datasets almost certainly include individuals not falling into these two categories (Lanfranco *et al.* 2004)). We make no statements in relation to gender, which is determined by many factors beyond genetics.

### Seventy-seven variants show genome-wide significance as candidates for sexually antagonistic selection

To identify autosomal variants that could be under sexually antagonistic selection, we performed a genome-wide association study (GWAS) between females and males in two large, independent cohorts (BioVU: 34,269 females and 27,491 males; UK Biobank: 264,813 females and 223,478 males). We first applied standard quality control steps to remove samples with high relatedness, discordant sex, or high heterozygosity and excluded genotyped variants with high overall missing rate (Methods). We account for potential confounders by including age and 12 principal components for population stratification as covariates. The resulting p-values are well-calibrated, as verified by permuting the sex labels in the UK Biobank cohort (Supplementary Figure 1A), and so the standard genome-wide significance threshold of P < 5E-8 is appropriate for the association analysis (Methods, Supplementary Figure 1B). Applying this threshold resulted in five and 72 genome-wide significant variants in BioVU and UK Biobank, respectively.

Since different amount of missing data for each variant between cases and controls can lead to spurious associations (Moskvina *et al.* 2006), we tested variants for a statistically significant difference in the missing rate between females and males (Methods). This control excluded what would have been 64 genome-wide significant variants in the UK Biobank and none in the BioVU cohort (Supplemental Figure 2, Supplementary File 1), leaving us with eight and five variants in the two datasets, respectively (Figure 1, Table 1). One intriguing genome-wide significant variant in the UK Biobank cohort, (rs11032483; OR = 1.25, P < 1.3E-53), which lies in a known regulatory region on chromosome 11 and has evidence from association studies for increasing risk in males and being protective in females for a number of sex-specific reproductive pathologies (Cortes *et al.* 2018).

**Table 1.**
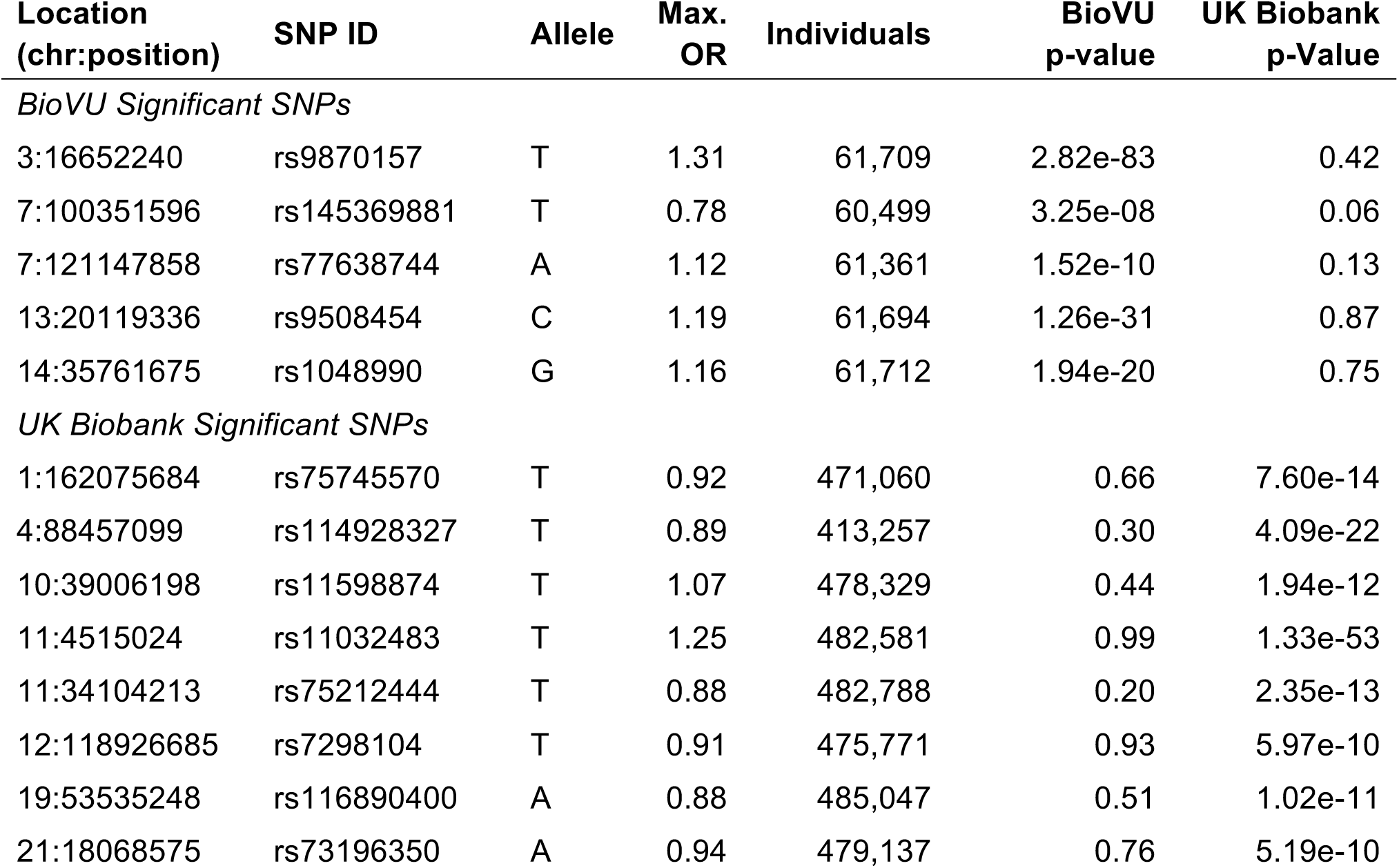
Genome-wide significant variants in BioVU and UK Biobank cohorts. Variants passing genome-wide significance (P < 5E-8) in the BioVU or UK Biobank cohorts are reported. Genome-wide significant variants did not replicate across the cohorts. Location is reported in GRch37/hg19 coordinates. Allele refers to the effect allele with which odds ratio (OR) is calculated. Individuals refers to the total number of individuals tested for the variant.

**Figure 1.**
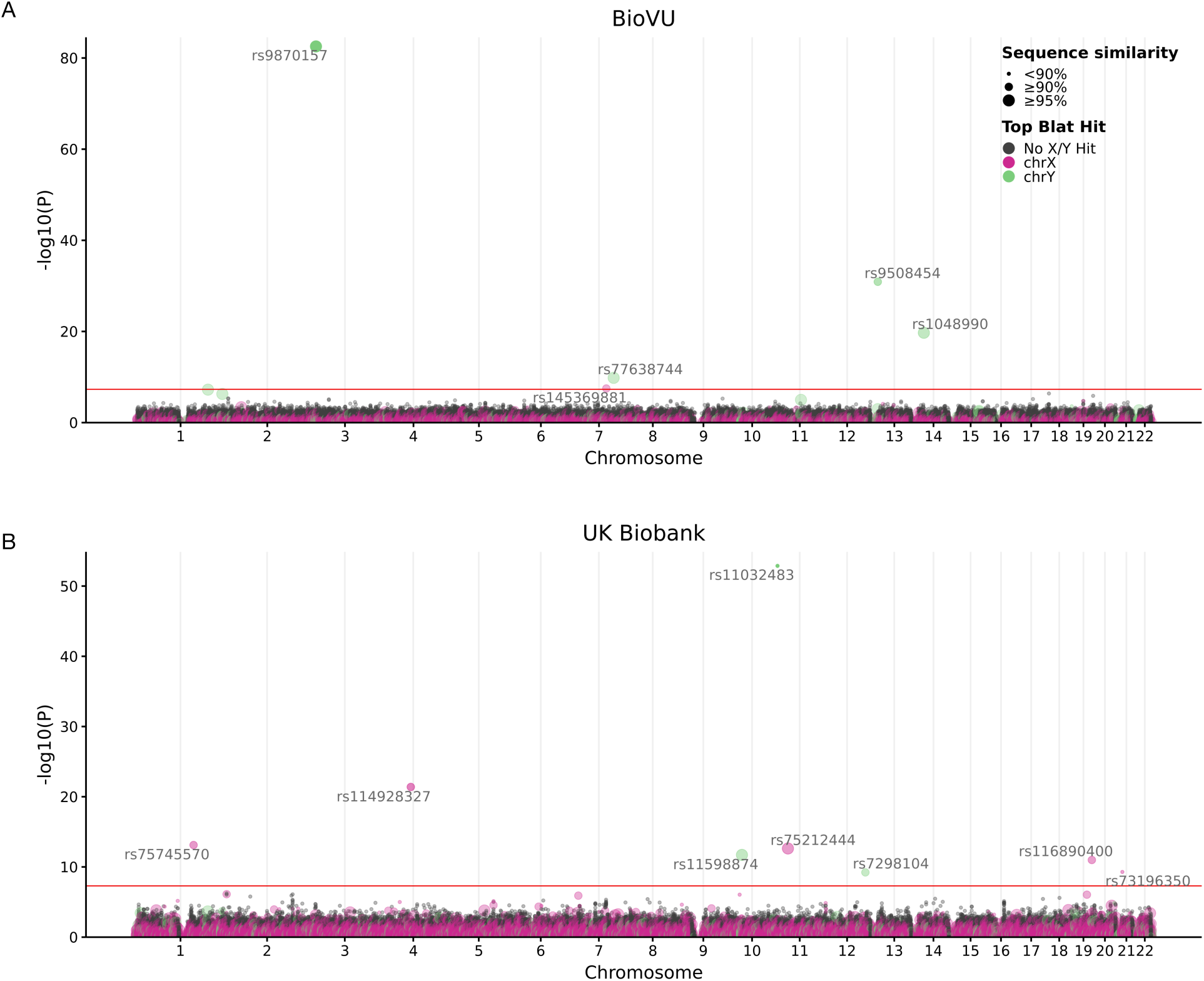
Genome-wide association tests for genetic sex reveals candidate variants for sexually antagonistic selection. To identify candidate variants for sexually antagonistic selection, we performed genome-wide association tests between females (cases) and males (controls) in two large biobank cohorts: **(A)** BioVU (females = 34,269, males = 27,491) and **(B)** UK Biobank (females = 264,813, males = 223,478). After standard quality control and sex-specific missingness filters (Methods), we identified five variants with genome-wide statistically significant associations (P < 5E-8, solid red line) in BioVU and eight in the UK Biobank. None of the significant variants in BioVU and UK Biobank replicated at genome-wide or nominal significance (P < 0.05) across the two cohorts (Table 1). The probe sequence for each associated variant (except rs11032483) had >90% sequence identity to at least one sequence on a sex chromosome (Table 2). Each point represents one variant. Each variant is colored by whether the best match of its probe sequence to a sex chromosome (according to BLAT score) is on X (pink) or Y (green). If it has no strong match to either sex chromosome it is colored black. The size of each point indicates the degree of sequence similarity.

**Figure 2.**
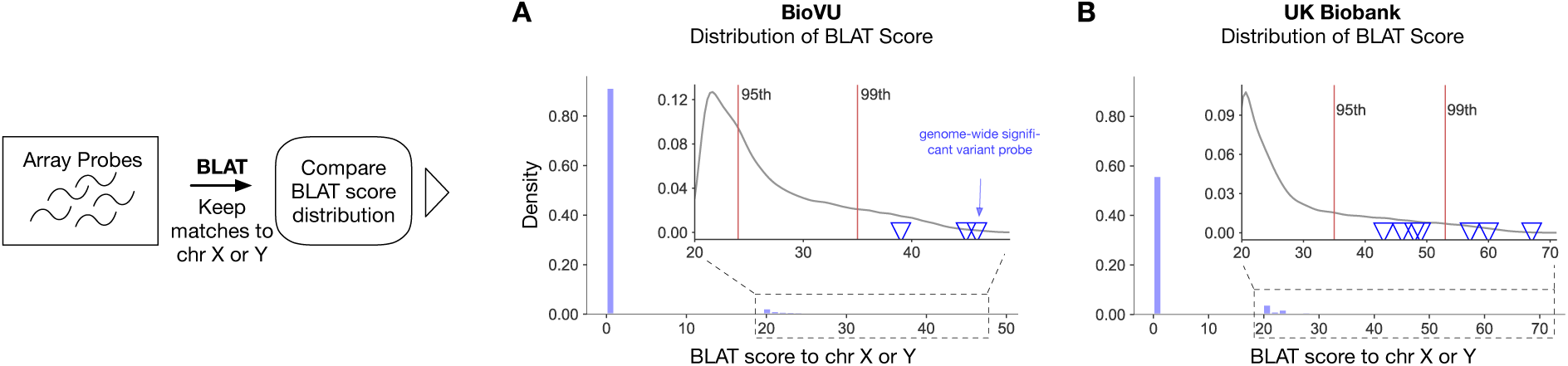
Probes for autosomal variants associated with genetic sex show high sequence similarity to sex chromosomes. We searched probe sequences used to genotype autosomal variants in the BioVU (798,051 autosomal probes) and UK Biobank (620,040 autosomal probes) cohorts for high sequence similarity to sex chromosome regions using BLAT (Methods). (**A)** More than 80% of BioVU autosomal probes do not have any sequence similarity (BLAT score ≤ 20) to a sex chromosome region; these are plotted at 0. Among the 83,083 BioVU probes with similarity to a sex chromosome sequence (inset), the probes for the variants with genome-wide significant associations with sex (blue triangles) are all in the tail of the distribution beyond the 99^th^ percentile of the BLAT match score. **(B)** Patterns are similar for the UK Biobank probes; however, a higher fraction (20%, 128,090) have detectable similarity to a sex chromosome, likely due to their greater length than the BioVU probes.

### No candidate loci replicate across BioVU and the UK Biobank

Comparing the five autosomal significant hits from BioVU to the eight from the UK Biobank, none of the associations are genome-wide significant in both cohorts (Table 1). Furthermore, none of the significant hits in one cohort even meet a nominal significance threshold (P < 0.05) in the other cohort. For example, the variant with the strongest association in the UK Biobank cohort (rs11032483) had no evidence for association with sex in the BioVU cohort (P = 0.99).

**Table 2.** Best sex chromosome sequence match for genome-wide significant variant probes. Variants with genome-wide significant associations with genetic sex are reported with GWAS p-value (P-value) and the matched sex chromosome region (Matched Sex Chromosome) with the highest BLAT score (BLAT score). The sequence similarity and length of the matching region (Match Length) are also reported.

| Dataset | SNP ID | Location (chr:position) | P-value | Matched Sex Chromosome | BLAT Score | Sequence Similarity (%) | Match Length (bp) |
| --- | --- | --- | --- | --- | --- | --- | --- |
| BioVU | rs9870157 | 3: 16652240 | 2.82E-83 | Y: 26964471-26964521 | 46 | 96.0 | 50 |
| BioVU | rs145369881 | 7: 100351596 | 3.25E-8 | X: 26864979-2685116 | 39 | 90.0 | 50 |
| BioVU | rs77638744 | 7: 121147858 | 1.52E-10 | Y: 23315613-23316169 | 45 | 98.0 | 50 |
| BioVU | rs9508454 | 13: 20119336 | 1.26E-31 | Y: 28612640-28612689 | 46 | 91.9 | 50 |
| BioVU | rs1048990 | 14: 35761675 | 1.94E-20 | Y: 15398460-15398510 | 46 | 96.0 | 50 |
| UKBB | rs75745570 | 1: 162075684 | 7.60E-14 | X: 121952043-121952114 | 57 | 90.2 | 71 |
| UKBB | rs114928327 | 4: 88457099 | 4.09E-22 | X: 79084149-79084223 | 60 | 93.0 | 71 |
| UKBB | rs11598874 | 10: 39006198 | 1.94E-12 | Y: 13568059-13568130 | 67 | 97.2 | 71 |
| UKBB | rs11032483 | 11: 4515024 | 1.33E-53 | Y: 19070733-19070803 | 48 | 84.3 | 70 |
| UKBB | rs75212444 | 11: 34104213 | 2.35E-13 | X: 36967486-36967536 | 46 | 96.0 | 50 |
| UKBB | rs7298104 | 12: 118926685 | 5.97E-10 | Y: 1513524-1513899 | 43 | 93.9 | 59 |
| UKBB | rs116890400 | 19: 53535248 | 1.02E-11 | X: 38605892-38605963 | 57 | 90.2 | 71 |
| UKBB | rs73196350 | 21: 18068575 | 5.19E-10 | X: 80780038-80780101 | 49 | 88.9 | 63 |

The regions surrounding each of the significant variants do not exhibit the expected association signal clusters arising from variants in strong linkage disequilibrium (LD) with the causal variant. For example, the most strongly associated variant overall (rs9870157) has 33 variants with R^2^ of at least 0.8 in the 1000 Genomes Phase 3 European-ancestry (EUR) populations. However, there are no other strong associations among these variants. The lack of replication across the two cohorts and the missing association peaks among variants in strong LD suggest that these signals could be false positives driven by technical or biological artifacts.

### Significant associations are likely due to mis-hybridization with sex chromosome regions

Genotyping error can occur due to probe cross-reactivity between different regions of the genome. Sex-biased error has been observed in array-based studies of DNA methylation (Chen *et al.* 2013) and has been reported in the canid genome (Tsai *et al.* 2019), the stickleback genome (Bissegger *et al.* 2019), and on the Y chromosome in humans (Boraska *et al.* 2012; Lucotte *et al.* 2016). For instance, if an autosomal variant is assayed with a probe sequence that has sufficient sequence similarity to a Y chromosome region carrying the reference allele, then males homozygous for the alternate allele at the autosomal locus may instead be genotyped as heterozygous for the alternate allele. Females would not be subject to this bias, and thus there would appear to be an allele frequency difference between the sexes. Similarly, an autosomal variant with a probe sequence with high similarity to the X chromosome could result in a lack of homozygotes for the allele not on the X chromosome in both sexes, but the strength of this effect would differ between females and males. Furthermore, such cross-reactivity can lead the normalized intensities produced by genotyping arrays to lie outside of the regions corresponding to each genotype, and thus a missing genotype (Zhao *et al.* 2018). Cross reactivity to a sex chromosome could therefore cause a differential missingness rate between the sexes. Indeed, we observe an almost complete lack of minor allele homozygotes in males across all thirteen genome-wide significant SNPs, as well as for females in all but four genome-wide significant SNPs (Supplemental Table 1). The same explanation is likely behind the 64 SNPs discarded for association between missingness and sex, as 26 of these SNPs have almost no male minor allele homozygotes and 47 have a p-value for lack of minor allele homozygotes of less than 1E-6.

To quantify the potential for mis-hybridization of sex chromosome regions to autosomal probes, we used BLAT (Kent 2002) to find all regions across the genome with high sequence similarity to autosomal probe sequences on the MEGAEx (BioVU) and UK Axiom/BilEVE (UK Biobank) genotyping arrays (Methods). We assign each probe sequence to the sex chromosome region with the highest BLAT score.

The probes for each significantly associated variant have high sequence similarity to a sex chromosome region (Figure 2, Table 2). In contrast, the majority of probes (79% in UK Biobank, 89% in BioVU) do not have any detectable similarity (BLAT score < 20) to a sex chromosome sequence. Compared to the distribution of BLAT scores for probes with a match to a sex chromosome region, all genome-wide significant variants had BLAT scores greater than the 99th and 95th percentile for BioVU and UK Biobank respectively (inset Figure 2A, 2B). Using a stricter criteria to define potential sex chromosome sequence similarity (Methods), we find that all genome-wide significant variants in BioVU (Supplemental Figure 3A) and six out of eight genome-wide significant variants in UK Biobank (Supplemental Figure 3B) still have strong sequence similarity to a sex chromosome region (Table 2). Only 0.57% (4,587 probes) and 3.3% (20,528) of all probes in BioVU and UK Biobank respectively have such a sex chromosome match (Supplemental Figure 3). The difference in percentage is likely due to the UK Biobank arrays having longer probe sequences. Probes of genome-wide significant variants have similar BLAT matching properties as non-significant probes (Supplementary Figure 4) in UK Biobank and BioVU. Many of the 64 SNPs discarded for between-sex differences in missingness also demonstrated high sequence similarity to sex chromosome regions (Supplementary Table 8). Overall, the lack of homozygotes and the high sequence similarity between significant probes and sex chromosomes strongly suggests that sex-specific genotyping error is the source of the significant associations rather than sexually antagonistic selection.

**Figure 3.**
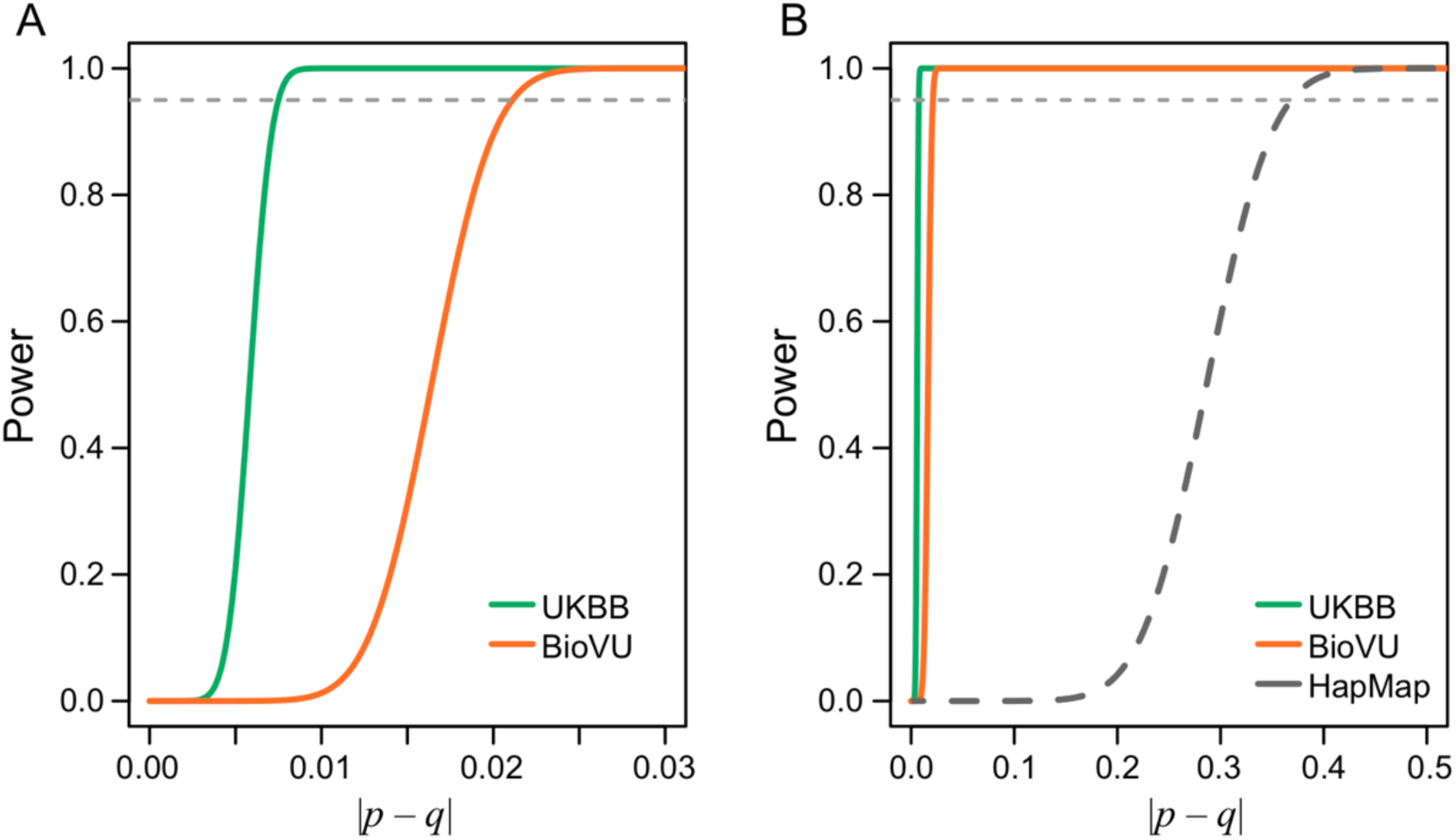
Statistical power was sufficient to detect small allelic divergence between the sexes. **(A)** The power to detect different levels of allelic divergence between the sexes was calculated for the BioVU (blue) and UKBB (green) cohorts. The dashed line shows the 95% power threshold. **(B)** Statistical power for the analyzed cohorts compared to previous analysis of human sequences (Lucotte *et al.* 2016) based on approximately 100 individuals per HapMap population (gray).

### The lack of sex-specific allele frequency differences is not due to being statistically underpowered

To determine if the lack of significant associations might be a result of being underpowered to detect plausible effect sizes, we conducted a power analysis (Methods). Based on the large cohort sizes, we have 95% power to detect a variant with a true allele frequency difference greater than 2% between the sexes in the BioVU cohort and greater than 0.8% in the UK Biobank (Figure 3A). A frequency difference of *f*% caused by sex-specific antagonistic selection at a locus requires a mortality of roughly *f/2*%, so we should be able to detect segregating variants with sex-specific mortality effects of at least 0.4% (Supplemental File 9). For comparison, a cohort of 100 individuals, as used in a previous HapMap study (Lucotte *et al.* 2016), only has 95% power to detect allele frequency differences between the sexes of 38% or greater (Figure 3B).

## DISCUSSION

Understanding how sex-specific effects are transmitted by autosomal variation is critical for understanding how sexual dimorphisms arise and fix in populations. Sexually antagonistic selection maintains sexual dimorphisms and is predicted to be a pervasive driver of genome evolution (Rowe *et al.* 2018). Yet, empirically, the genomic signature of this process is not well characterized. In this study, we sought to identify the extent of one genomic signature of sexually antagonistic viability selection acting on autosomal variation in human populations. Capitalizing on two of the largest available biobanks, we performed genome-wide association tests for genetic sex that failed to identify and replicate any genome-wide significant variants. On closer inspection, a number of promising genome-wide significant variants were driven by technical artifacts, most likely due to high sequence similarity to a sex chromosome. We conclude there is no conclusive signal in these data of sexually antagonistic viability selection acting on genetic variants at individual loci based on a male-female divergence statistic.

These results stand in contrast to recent male-female *F*_*ST*_ studies, that have reported tens to hundreds of significantly differentiated variants (Lucotte *et al.* 2016; Flanagan and Jones 2017; Wright *et al.* 2018; Dutoit *et al.* 2018; Bissegger *et al.* 2019; Vaux *et al.* 2019). These studies suggest strong, pervasive sexually antagonistic viability selection acting across the genomes of various species, which would be puzzling in light of theoretical observations and simulations indicating that strong allelic divergence between the sexes requires high sex-specific mortality rates to overcome the homogenizing effect of meiotic segregation occurring every generation (Kasimatis *et al.* 2019). In contrast to these studies, the sample size of our study provided statistical power to distinguish true signal of plausible magnitude from stochastic noise. Additionally, our use of larger sample sizes provided power to detect smaller allelic divergence between the sexes – within the range predicted to be generated by weak sexually antagonistic selection. Our results are in line with a previous meta-analysis of sex-specific common variant differences in humans (Boraska *et al.* 2012), though our direct, replicated approach with larger sample sizes mitigates against potential confounders across different studies.

We found strict quality control measures for population structure and multiple testing essential. In particular, rigorous testing for sequence similarity to the sex chromosomes showed that all significant SNPs had strong sequence matches. The potential for high sequence similarity between autosomes and sex chromosomes to generate sex-biased genotyping errors has been reported previously (Chen *et al.* 2012; 2013). However, the potential for these sex chromosome artifacts to affect population genetic statistics has not been fully appreciated until recently (Bissegger *et al.* 2019; Tsai *et al.* 2019) or has only been examined for the Y chromosome (Lucotte *et al.* 2016). In particular, probe sequences with high sequence similarity to one the sex chromosomes can lead to skewed allele frequency estimates in a sex-specific manner due to sequence mis-hybridization and the different sex chromosome content between females and males. This problem extends beyond SNP-based genotyping to read-based sequencing data, where inaccurate mapping of reads to an autosome instead of the sex chromosome could generate a similar skew in allele frequencies. This sex chromosome effect is potentially very common, and therefore, must be explicitly considered in any sex-specific or sex-stratified analyses to prevent technical and bioinformatic artifacts from generating false signals. Participation bias rather than differential mortality can also generate a signal of male-female divergence (Pirastu *et al.* 2020), though this source of error is not relevant in this study since we did not find candidate SNPs for sexually antagonistic selection that passed our quality controls. Such artifacts will be especially problematic in species with new sex chromosomes, poorly assembled genomes, or rapidly evolving sex chromosome systems. In our case, filtering out SNPs with large differences in missingness between sexes and/or lack of homozygotes was sufficient to remove problematic SNPs.

Comparison of sequence similarity and match length for all probes indicates that thousands of other probes have similarly strong sex chromosome matches as the candidate variants analyzed here (Supplementary Figure 2). While previous studies have detected similar hybridization effects (Chen *et al.* 2012; 2013), the extent to which they can skew association results has not yet been reported on the UKBiobank and BioVU arrays. This high sequence similarity could suggest that more variants should show false positive signatures of sex-specific allele frequency differences. However, multiple factors contribute to the potential for mis-hybridization and inaccurate genotyping. For example, hybridization strength and kinetics are determined by sequence attributes beyond simple sequence identity, including local GC content and the potential for DNA secondary structures to form (Zhang *et al.* 2018). Furthermore, the sequence region matched on the sex chromosome (i.e., pseudo-autosomal versus non-recombining) also matters. It is also likely that different quality control strategies used on different genotyping array platforms filter different problematic sites.

Although sexually antagonistic selection is certainly an important selective pressure, we see no evidence of it generating substantial autosomal allelic divergence between the sexes in human populations. This strong negative result is unusual, as genome-wide association studies for most traits on a biobank-scale find significantly associated SNPs. We know that humans have the opportunity for sexually antagonistic effects, as seen through sex-specific mortality and disease susceptibility (Morrow 2015; Khramtsova *et al.* 2018). However, randomization of alleles every generation by meiotic segregation means that a large selective pressure is required to create a large difference in allele frequencies, and thus, this genetic process makes it harder to detect the results of sexually antagonistic selection. Furthermore, some sexually antagonistic variants are not stably polymorphic; we would not detect these because they move rapidly to fixation (Rowe *et al.* 2018; Kasimatis *et al.* 2019).

Given the confounding factors, technical artifacts, and high sampling variance, identifying variants with small sex-specific effect sizes is a formidable challenge. We strongly recommend that future studies avoid simple metrics, like the male-female *F*_*ST*_, and instead incorporate strict quality filters and control for known confounders into their association tests. Sexually antagonistic viability selection is not the only action of sex-specific selection nor is male-female allelic divergence at a single locus the only possible signature of sexual antagonism. Given the extent of sexual dimorphisms in nature, there are almost surely autosomal loci subject to sexually antagonistic selection, which may be detectable through other genomic signatures. However, our work illustrates that the field must reconsider our assumptions and develop new metrics for identifying signatures of sexual antagonism in the light of theoretical expectations to understand how this process affects the genome. Such studies will help us understand the translation of sex across the genotype-phenotype map and apply this to human health.

## Supporting information

Kasimatis_Abraham_Supplements

## Acknowledgements

We thank Locke Rowe and the Phillips and Capra lab groups for their comments on this work, as well as Clare Bycroft, Adrián Cortés, and Gil McVean for information about UK Biobank genotyping. A.K. was supported by NIH award R01GM117241. J.A.C. was supported by NIH award R35GM127087. P.C.P. was supported by NIH award R01GM102511 and R35GM131838. A.A. is supported by NIGMS of the National Institutes of Health under award number T32GM007347. This research has been conducted using the UK Biobank Resource under Application Number 43626. This work was conducted in part using the resources of the Advanced Computing Center for Research and Education at Vanderbilt University and the resources of the Research Advanced Computing Services at the University of Oregon. The content is solely the responsibility of the authors and does not necessarily represent the official views of the National Institutes of Health.

## Contributions

K.R.K. and P.C.P. devised the project. K.R.K., A.A., P.L.R., A.D.K., J.A.C., and P.C.P. designed the analyses. K.R.K. and A.A. performed analyses. K.R.K., A.A., P.L.R, and J.A.C wrote the manuscript with the support of the other authors.

The authors declare no competing interests.

## SUPPLEMENTARY FILES

**Supplementary File 1:** “TableS1_gwas_significant_hits.xlsx” - Summary statistics for genome-wide significant variants (including those removed for uneven missing rate between the sexes) associated with genetic sex in BioVU and UK Biobank. MALE_HOM1, MALE_HET, and MALE_HOM2 are the counts of genotypes of the minor allele homozygote, heterozygote, and major allele homozygote genotype calls in males, respectively; MALE_MISSING is the number of missing genotypes in males reported by plink. FEM_ prefixes similar columns for females. MISSING_PVAL gives the p-value from Fisher’s exact test comparing proportions of missing genotypes between males and females as reported by plink. HOM1_PVAL gives the p-value for a binomial test for the proportion of minor allele homozygotes (of either sex) being equal to the marginal allele frequency squared. OR, STAT and ASSOC_PVAL gives the maximum odds ratio, t-statistic and p-value from logistic regression as described in the text.

**Supplementary File 2:** “TableS2_HWE_genotype_counts.xlsx” - Genome-wide significant variants associated with genetic sex in BioVU and UK Biobank with Hardy-Weinberg Equilibrium statistics.

**Supplementary File 3:** “TableS3_raw_blat_xy_hits_bv.tsv” - Raw BLAT results for hits to chromosome X or Y for probes in the MEGAex genotyping array.

**Supplementary File 4:** “TableS4_raw_blat_xy_hits_ukaxiom.tsv” - Raw BLAT results for hits to chromosome X or Y for probes in the UK Biobank Axiom genotyping array.

**Supplementary File 5:** “TableS5_raw_blat_xy_hits_ukbil.tsv” - Raw BLAT results for hits to chromosome X or Y for probes in the UK BilEVE genotyping array.

**Supplementary File 6:** “TableS6_best_blatscore_xy_hit_length_filtered_bv_gwas.tsv” - Best BLAT matches to chromosome X or Y based on highest BLAT score for each probe in the MEGAex genotyping array.

**Supplementary File 7:** “TableS7_best_blatscore_xy_hit_length_filtered_uk_gwas.tsv” - Best BLAT matches to chromosome X or Y based on highest BLAT score for each probe in the UK Biobank. BLAT results for hits to chromosome X or Y for probes are chosen after pooling across UK Biobank Axiom and UK BilEVE genotyping arrays.

**Supplementary File 8:** “TableS8_uk_var_w_missingness_best_blatscore_xy.tsv” – Best sex chromosome match based on highest BLAT score for UK Biobank genome-wide significant variants with statistically significant difference in missing rate between females and males.

**Supplementary File 9:** “FileS9_mortality_selection_derivation.pdf” – Mathematical derivation for estimating the sex-specific mortality cost.

**Supplementary Figure 1.**
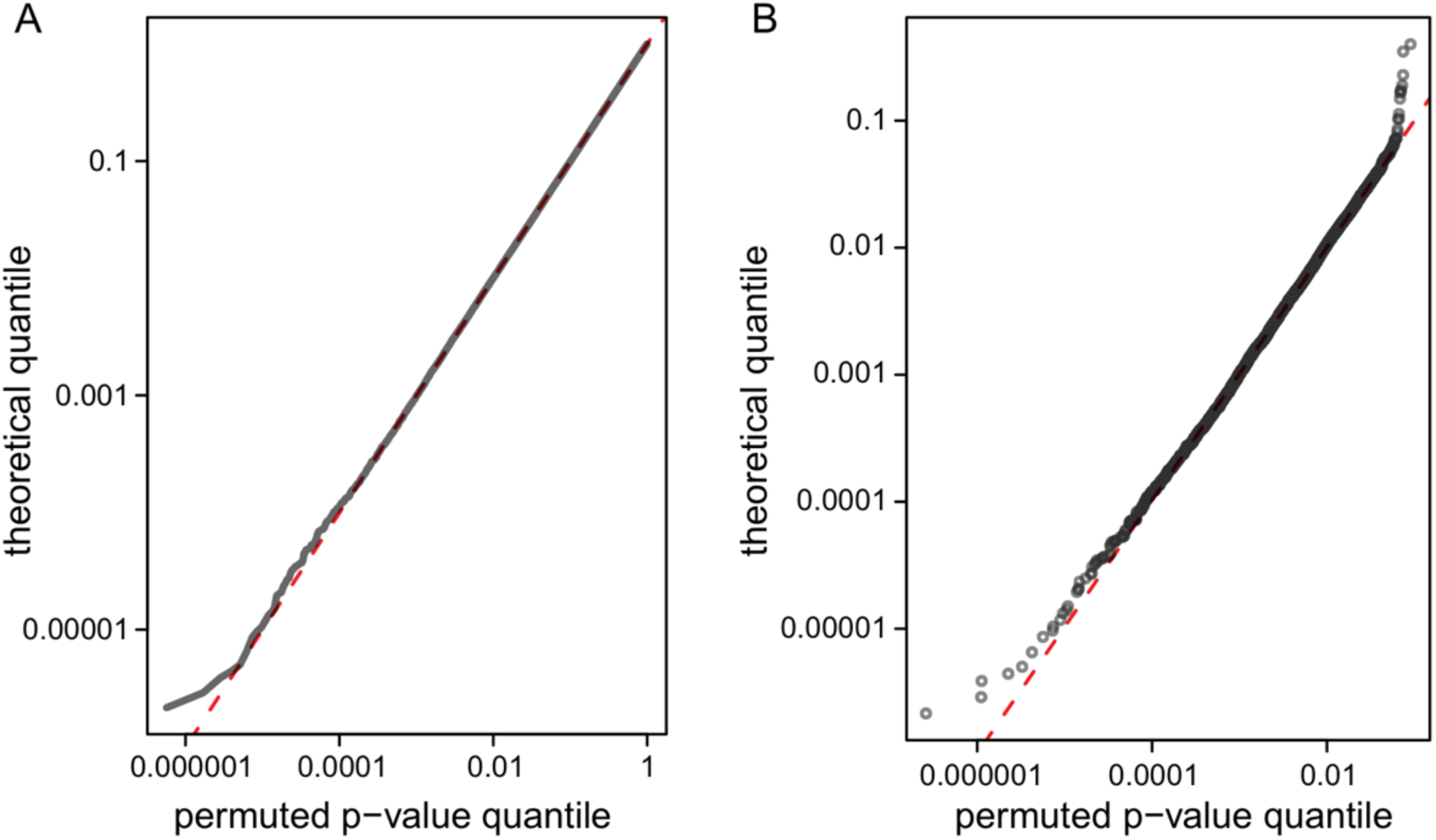
Permutation of genetic sex to generate a null distribution demonstrates that p-values are well calibrated. We randomly permuted genetic sex and ran a genome-wide association test between the permuted females and males in the UKBB cohort 100 times. Only those variants with a p-value < 0.01 under the association with the true genetic sex are considered (n = 8,868 SNPs). **(A)** Q-Q plot of all the permuted variants shows they are uniformly distributed. **(B)** Permuted variants are uniformly distributed even at very small p-values.

**Supplementary Figure 2.**
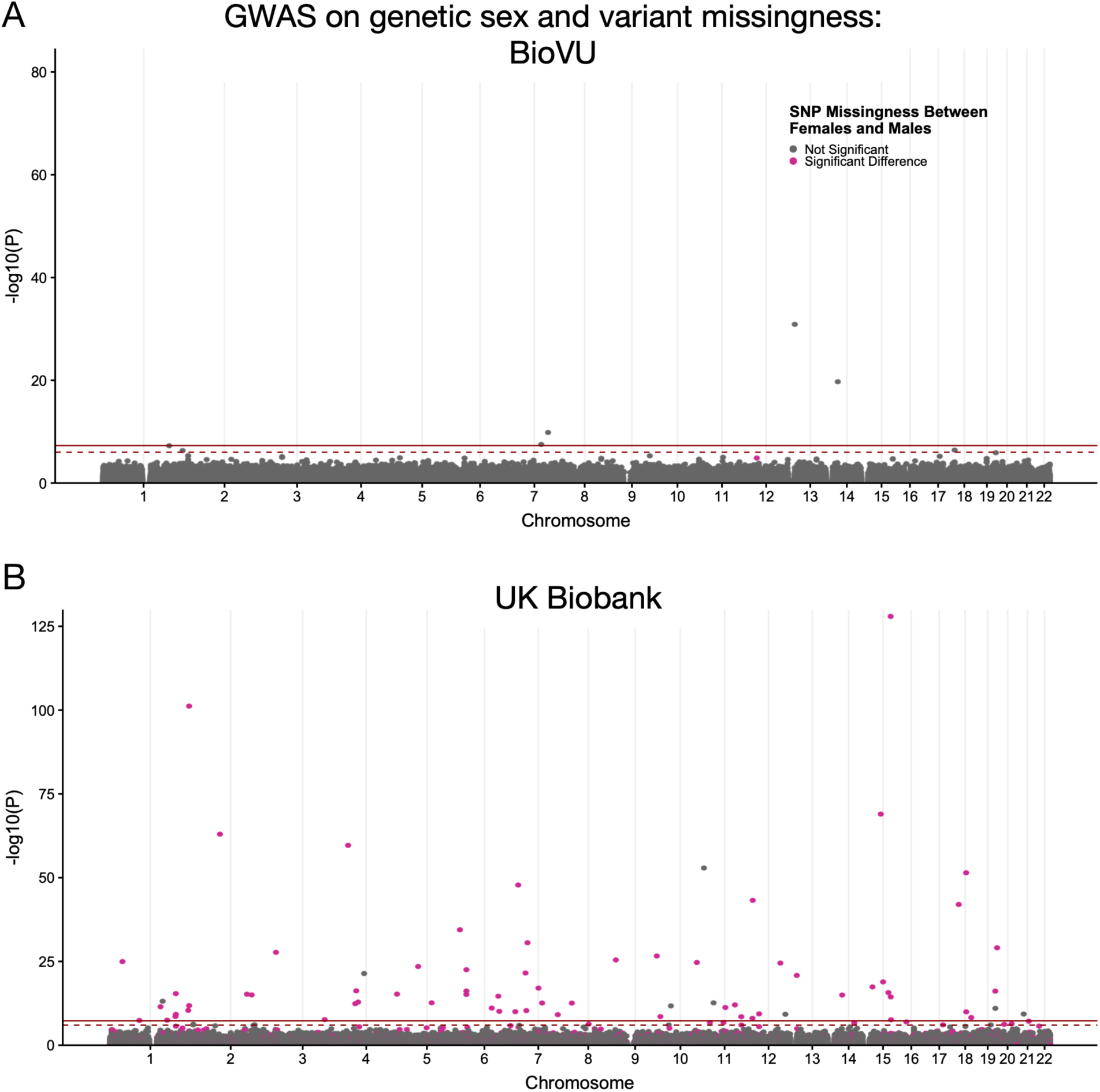
Significantly different variant missingness between females and males contribute many spurious association in the UK Biobank GWAS for genetic sex. After running a GWAS for genetic sex in **(A)** BioVU and **(B)** UK Biobank cohorts, we identify five and 72 variants with genome-wide significant associations (solid red line, P < 5E-8; dashed red line P < 5E-6) respectively. Variants with a statistically significant difference (p < 0.00001, Fisher’s Exact test) in the missing rate between females and males are colored in red. In the UK Biobank cohort, 64 genome-wide significant variants also have a statistically significant difference in the missingness between cases and control, suggesting that these associations are spurious.

**Supplementary Figure 3.**
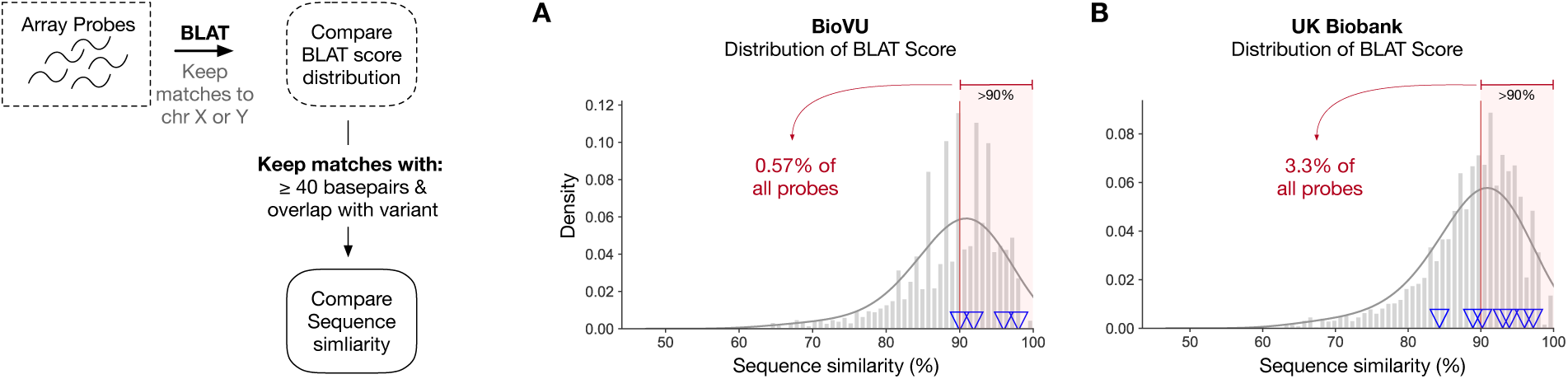
Sequence similarity distribution of probes after applying strict matching criteria to a sex chromosome. To identify probes most likely to mis-hybridize between autosomal and sex chromosome sequences, we filtered those whose best BLAT match met the following criteria in (A) BioVU and (B) UK Biobank: sex chromosome match ≥40 base pairs in length, ≥90% sequence similarity, and overlap of the matching region with the genotyped variant. Out of all autosomal probes, 0.57% and 3.3% met the aforementioned criteria in BioVU and UK Biobank, respectively.

**Supplementary Figure 4.**
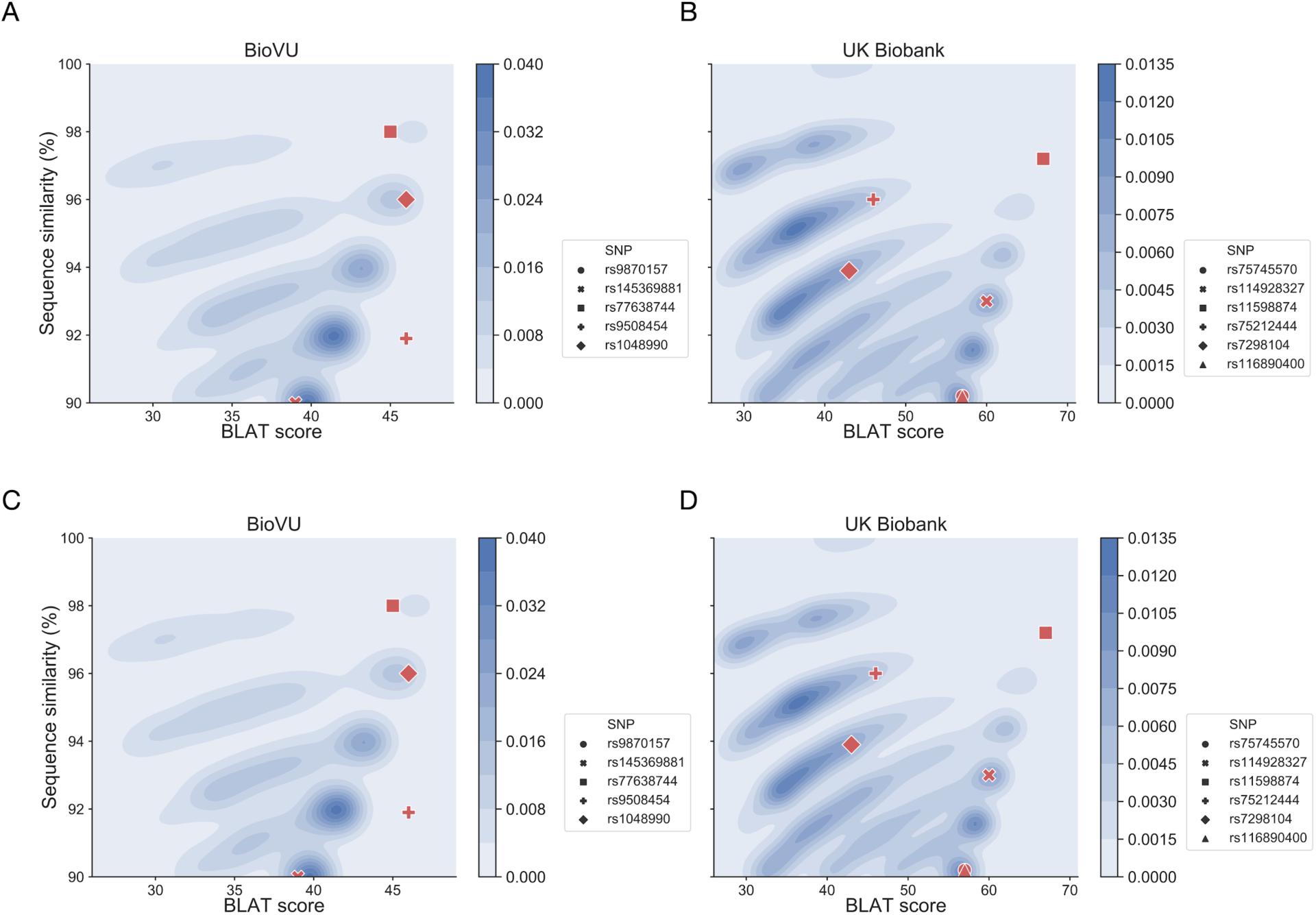
Probes of genome-wide significant variants with a match to sex chromosome regions have similar matching properties as non-significant variant when comparing BLAT score and match length to sequence similarity. Using BLAT, we identify array probe sequences with high sequence similarity (≥90%) to a sex chromosome region, have a match length ≥40 base pairs, and overlap or is adjacent on the probe sequence to the variant being genotyped. We plot bivariate kernel density estimates comparing **(A)** BLAT score and **(B)** match length against sequence similarity (y-axis) for BioVU and UK Biobank probe sequences. Darker blue represents areas of higher density. The position of probe sequences for genome-wide significant variants are overlaid as red markers on each plot. Comparing against the densities of the non-significant variants, probes of genome-wide significant variants occur in areas of high density suggesting they have similar matching properties as non-significant probes.

## Notes

### Competing Interest Statement

The authors have declared no competing interest.

## REFERENCES

Archer C. R., Recker M., Duffy E., Hosken D. J., 2018 Intralocus sexual conflict can resolve the male-female health-survival paradox. Nat. Commun. 9: 1–7.

Arnqvist G., Rowe L., 2005 Sexual conflict. Princeton University Press, Princeton, NY.

Bissegger M., Laurentino T. G., Roesti M., Berner D., 2019 Widespread intersex differentiation across the stickleback genome - The signature of sexually antagonistic selection? Mol. Ecol. 77: 1–10.

Boraska V., Jeroncic A., Colonna V., Southam L., Nyholt D. R., et. al., 2012 Genome-wide meta-analysis of common variant differences between men and women. Hum. Mol. Genet. 21: 4805–4815.

Bycroft C., Freeman C., Petkova D., Band G., Elliott L. T., et. al., 2018 The UK Biobank resource with deep phenotyping and genomic data. Nature 562: 203–209.

Chang C. C., Chow C. C., Tellier L. C., Vattikuti S., Purcell S. M., et. al., 2015 Second-generation PLINK: rising to the challenge of larger and richer datasets. Gigascience 4: 2–16.

Chen Y.-A., Choufani S., Grafodatskaya D., Butcher D. T., Ferreira J. C., et. al., 2012 Cross-Reactive DNA Microarray Probes Lead to False Discovery of Autosomal Sex-Associated DNA Methylation. Am. J. Hum. Genet. 91: 762–764.

Chen Y.-A., Lemire M., Choufani S., Butcher D. T., Grafodatskaya D., et. al., 2013 Discovery of cross-reactive probes and polymorphic CpGs in the Illumina Infinium HumanMethylation450 microarray. Epigenetics 8: 203–209.

Cheng C., Kirkpatrick M., 2016 Sex-specific selection and sex-biased gene expression in humans and flies. PLoS Genetics 12: e1006170.

Cortes A., Dendrou C. A., Fugger L., McVean G., 2018 Systematic classification of shared components of genetic risk for common human diseases. bioRxiv: 1–22.

Cox R. M., Calsbeek R., 2009 Sexually antagonistic selection, sexual dimorphism, and the resolution of intralocus sexual conflict. Am. Na.t 173: 176–187.

Das S., Forer L., Schönherr S., Sidore C., Locke A. E., et. al., 2016 Next-generation genotype imputation service and methods. Nat. Genet. 48: 1284–1287.

Deegan D. F., Engel N., 2019 Sexual dimorphism in the age of genomics: how, when, where. Front. Cell. Dev. Biol. 7: 1–7.

Dumitrescu L., Barnes L. L., Thambisetty M., Beecham G., Kunkle B., et. al., 2019 Sex differences in the genetic predictors of Alzheimer’s pathology. Brain 142: 2581–2589.

Dutoit L., Mugal C. F., Bolívar P., Wang M., Nadachowska-Brzyska K., et. al., 2018 Sex-biased gene expression, sexual antagonism and levels of genetic diversity in the collared flycatcher (*Ficedula albicollis*) genome. Mol. Ecol. 27: 3572–3581.

Farquhar C. M., Bhattacharya S., Repping S., Mastenbroek S., Kamath M. S., et. al., 2019 Female subfertility. Nat. Rev. Dis. Primers. 5: 1–22.

Ferretti M. T., Iulita M. F., Cavedo E., Chiesa P. A., Schumacher Dimech A., et. al., 2018 Sex differences in Alzheimer disease - the gateway to precision medicine. Nat. Rev. Neurol. 14: 457–469.

Flanagan S. P., Jones A. G., 2017 Genome-wide selection components analysis in a fish with male pregnancy. Evol. 71: 1096–1105.

Kasimatis K. R., Nelson T. C., Phillips P. C., 2017 Genomic signatures of sexual conflict. J. Hered. 108: 780–790.

Kasimatis K. R., Ralph P. L., Phillips P. C., 2019 Limits to genomic divergence under sexually antagonistic selection. G3 9: 3813–3824.

Kent W. J., 2002 BLAT - The BLAST-like alignment tool. Genome Res. 12: 656–664.

Khramtsova E. A., Davis L. K., Stranger B. E., 2018 The role of sex in the genomics of human complex traits. Nat. Rev. Genet 62: 1–190.

Lanfranco F., Kamischke A., Zitzmann M., Nieschlag E., 2004 Klinefelter’s syndrome. Lancet 364: 273–283.

Lucotte E. A., Laurent R., Heyer E., Ségurel L., Toupance B., 2016 Detection of allelic frequency differences between the sexes in humans: a signature of sexually antagonistic selection. Genome Biol. Evo.l 8: 1489–1500.

Mank J. E., 2017 Population genetics of sexual conflict in the genomic era. Nat. Rev. Genet. 7: 1–10.

Morrow E. H., 2015 The evolution of sex differences in disease. Biol. Sex Differ. 6: 1–7.

Moskvina V., Craddock N., Holmans P., Owen M. J., O’Donovan M. C., 2006 Effects of differential genotyping error rate on the type I error probability of case-control studies. Hum. Hered. 61: 55–64.

Pirastu N., Cordioli M., Nandakumar P., Mignogna G., Abdellaoui A., et. al., 2020 Genetic analyses identify widespread sex-differential participation bias. bioRxiv: 1–54.

Rice W. R., Chippindale A. K., 2001 Intersexual ontogenetic conflict. J. Evol. Biol. 14: 685–693.

Rowe L., Chenoweth S. F., Agrawal A. F., 2018 The genomics of sexual conflict. Am. Nat. 192: 274–286.

Tsai K. L., Evans J. M., Noorai R. E., Starr-Moss A. N., Clark L. A., 2019 Novel Y chromosome retrocopies in canids revealed through a genome-wide association study for sex. Genes 10: 320–11.

Vaux F., Rasmuson L. K., Kautzi L. A., Rankin P. S., Blume M. T. O., et. al., 2019 Sex matters: otolith shape and genomic variation in deacon rockfish (*Sebastes diaconus*). Ecol Evol 27: 477–21.

Weedon M. N., Jackson L., Harrison J. W., Ruth K. S., Tyrrell J., et. al., 2019 Assessing the analytical validity of SNP-chips for detecting very rare pathogenic variants: implications for direct-to-consumer genetic testing. bioRxiv: 696799.

Wright A. E., Fumagalli M., Cooney C. R., Bloch N. I., Vieira F. G., et. al., 2018 Male-biased gene expression resolves sexual conflict through the evolution of sex-specific genetic architecture. Evol. Letters 215: 403–10.

Wright C. F., West B., Tuke M., Jones S. E., Patel K., et. al., 2019 Assessing the pathogenicity, penetrance, and expressivity of putative disease-causing variants in a population setting. Am. J. Hum. Genet. 104: 275–286.

Zhang J. X., Fang J. Z., Duan W., Wu L. R., Zhang A. W., et. al., 2018 Predicting DNA hybridization kinetics from sequence. Nat. Chem. 10: 91–98.

Zhao S., Jing W., Samuels D. C., Sheng Q., Shyr Y., et. al., 2018 Strategies for processing and quality control of Illumina genotyping arrays. Brief Bioinform. 19: 765–775.

